# Sex-Specific Cardiac Remodeling in Aged Rats after Early-Life Chronic Stress: Associations with Endocrine and Metabolic Factors

**DOI:** 10.1101/2024.04.03.587944

**Authors:** Carley Dearing, Ella Sanford, Nicolette Olmstead, Rachel Morano, Lawson Wulsin, Brent Myers

## Abstract

**Background:** Cardiovascular disease is a leading cause of death worldwide. Rates of cardiovascular disease vary both across the lifespan and between sexes. While multiple factors, including adverse life experiences, impact the development and progression of cardiovascular disease, the potential interactions of biological sex and stress history on the aged heart are unknown. To this end, we examined sex- and stress-specific impacts on left ventricular hypertrophy (VH) after aging. We hypothesized that early life chronic stress exposure impacts behavioral and physiologic responses that predict cardiac remodeling in a sex-specific manner.

**Methods:** Histological analysis was conducted on hearts of male and female rats previously exposed to chronic variable stress during the late adolescent period (postnatal days 43-62). These animals were challenged with a forced swim test and a glucose tolerance test before aging to 15 months and again being challenged. Predictive analyses were then used to isolate factors that relate to cardiac remodeling among these groups.

**Results:** Early-life chronic stress impacted cardiac remodeling in a sex-specific manner. Among rats with a history of chronic stress, females had increased inward VH. However, there were few associations within the female groups among individual behavioral and physiologic parameters and cardiac remodeling. While males as a group did not have VH after chronic stress, they exhibited multiple individual associations with cardiac susceptibility. Passive coping in young males and active coping in aged males related to VH in a stress history-dependent manner. Moreover, baseline corticosterone positively correlated with VH in unstressed males, while chronically-stressed males had positive correlations between VH and visceral adiposity.

**Conclusions:** These results indicate that females as a group are uniquely susceptible to the effects of early-life stress on cardiac remodeling later in life. Conversely, males have more individual differences in vulnerability, where susceptibility to cardiac remodeling relates to endocrine, metabolic, and behavioral measures depending on stress history. These results ultimately support a framework for accessing cardiovascular risk based on biological sex and prior adverse experiences.

**Highlights:**

- Aged female rats had greater left ventricular hypertrophy (VH) than males after early-life chronic variable stress.
- Tertile divisions based on susceptibility or resilience to inward VH indicated interactions between VH, sex, and stress on multiple behavioral and physiological measures.
- In males, VH correlated with endocrine and metabolic parameters in a stress history-dependent manner.
- Prior adverse experience and biological sex interact across the lifespan to impact cardiovascular risk.

**Plain English Summary:** Cardiovascular disease is the leading cause of death worldwide. Multiple factors influence the incidence and severity of cardiovascular disease including adverse life experiences, biological sex, and age. Alterations of heart structure predict negative cardiovascular health by impacting blood circulation; however, the potential interactions of stress history and biological sex on the aged heart are unknown. In this study, we examined how chronic stress exposure impacts heart structure in male and female rats after aging. Adolescent male and female rats were chronically stressed and then acutely challenged to examine behavioral, endocrine, and metabolic parameters both immediately following chronic stress and after aging. Heart morphology was quantified to examine how behavioral and physiological responses related to cardiac remodeling. Our results indicate that, as a group, female rats previously exposed to chronic stress were uniquely susceptible to inward remodeling of the heart. Subjects were further divided into sub-groups based on the level of inward remodeling of the ventricle. While male rats did not exhibit group effects on heart structure, individual variability in male heart morphology related to endocrine and metabolic parameters in a stress history-dependent manner. Here, there were interactions with multiple systems including coping behavior, stress hormones, and body composition. Moreover, males without a prior history of chronic stress had correlations between stress hormones and the degree of heart remodeling. However, males that were exposed to chronic stress had correlations between heart structure and abdominal fat. Overall, our results indicate that biological sex and stress history interact to predict cardiovascular susceptibility.

## 1. Background

Cardiovascular disease (CVD) contributes substantially to global disease burden [1] and is the leading cause of death in women [2]. Moreover, sex differences in the incidence of CVD vary across the lifespan [3]. While rates of CVD steadily increase with age in men, women generally experience lower CVD rates until menopause, at which point female CVD incidence increases to exceed that of men [4]. These differences are proposed to result from cardiovascular-protective effects of female sex steroids [5,6], centrally estradiol, that are lost with reproductive senescence [7,8]. However, aging-associated changes in physiology, including elevated blood pressure and glucose intolerance, impact the pathogenesis of CVD and associate with negative cardiovascular outcomes [9]. Ultimately, alterations in autonomic nervous system balance may underly the concentric cardiac hypertrophic remodeling that reduces ventricular volume and increases cardiac workload [10]. While left ventricular hypertrophy (VH) increases the risk of major cardiovascular events and heart failure [11], little is known about how early-life adversity and associated disruptions of autonomic-endocrine integration impact the cardiovascular system across the lifespan. Thus, differences in physiological stress response systems may account for sex-specific susceptibility to negative cardiovascular remodeling after aging.

Autonomic and neuroendocrine responses to stressors are essential for energy mobilization and adaptive homeostatic responses [12]. However, prolonged or repeated stressor exposure is associated with cardiometabolic dysfunction and neuropsychiatric disorders [13–15]. Exposure to stressors in early-life has immediate and long-term impacts on cognition and physiological stress reactivity in human and pre-clinical studies [16–18]. In particular, rodent studies of adolescent chronic stress have found persistent increases in male glucocorticoid stress reactivity [19,20]. Additionally, our prior work found that adolescent chronic variable stress (CVS) increases glucocorticoid responses to the forced swim test (FST) in both male and female rats. However, only female rats have impaired glucose clearance after CVS, which manifests as altered glucoregulation in later life [21]. Yet, the long-term consequences of early-life chronic stress for sex-specific cardiovascular health and disease susceptibility are unknown.

We previously generated a large cohort of male and female rats for a longitudinal study of adolescent CVS to examine both the immediate and persistent consequences for coping behavior and glucoregulation [21]. In the current study, we histologically examined cardiac structure in this cohort to test the hypothesis that sex-specific behavioral and endocrine measures predict cardiac susceptibility. To this end, we quantified concentric VH in animals that were exposed to adolescent CVS and then aged to 15 months. Paired assessments of behavioral coping and glucoregulation both immediately following CVS and at 15 months of age permitted correlational analyses of factors relating to VH susceptibility and resilience. These results ultimately shed light on the importance of biological sex and stress history for cardiac health and associations with behavioral, endocrine, and metabolic outcomes.

## 2. Methods

### 2.1 Subjects

Experiments were approved by the Institutional Animal Care and Use Committee of the University of Cincinnati (protocol 04-08-03-01) and complied with the National Institutes of Health Guidelines for the Care and Use of Laboratory Animals. All rats had daily welfare assessments by veterinary and/or medical service staff. Animals were kept on 12:12 light cycles with food and water available *ad libitum*. Sprague-Dawley rats from Envigo (Cumberalnd, VA) were bred in-house, producing 12 simultaneous litters of 10 rats each. Animals were weaned into same-sex group housing at PN24 and then pair-housed beginning PN35 through the remainder of the study. All estrous cycle phases were represented in female groups [21].

### 2.2 Design

Tissue was collected from all animals and hearts were histologically analyzed to examine relative VH. As previously reported [18], the experimental design involved randomly assigning 120 adolescent rats to CVS (n = 36/sex) and No CVS (n = 24/sex) groups with equal representation from each litter (n = 3/sex/litter CVS, n = 2/sex/litter No CVS) (Fig. 1A). These animals were exposed to the CVS paradigm [20,22] from PN 43-62 corresponding to the period of late adolescence [20]. All subjects then underwent acute psychogenic (FST) and metabolic (intraperitoneal glucose tolerance test [GTT]) challenges with blood sampled. Animals then aged to 15 months and received the FST and GTT again prior to tissue collection. Methodology and group data from the FST, GTT, and endocrine assessments were previously published [21].

**Fig 1.**
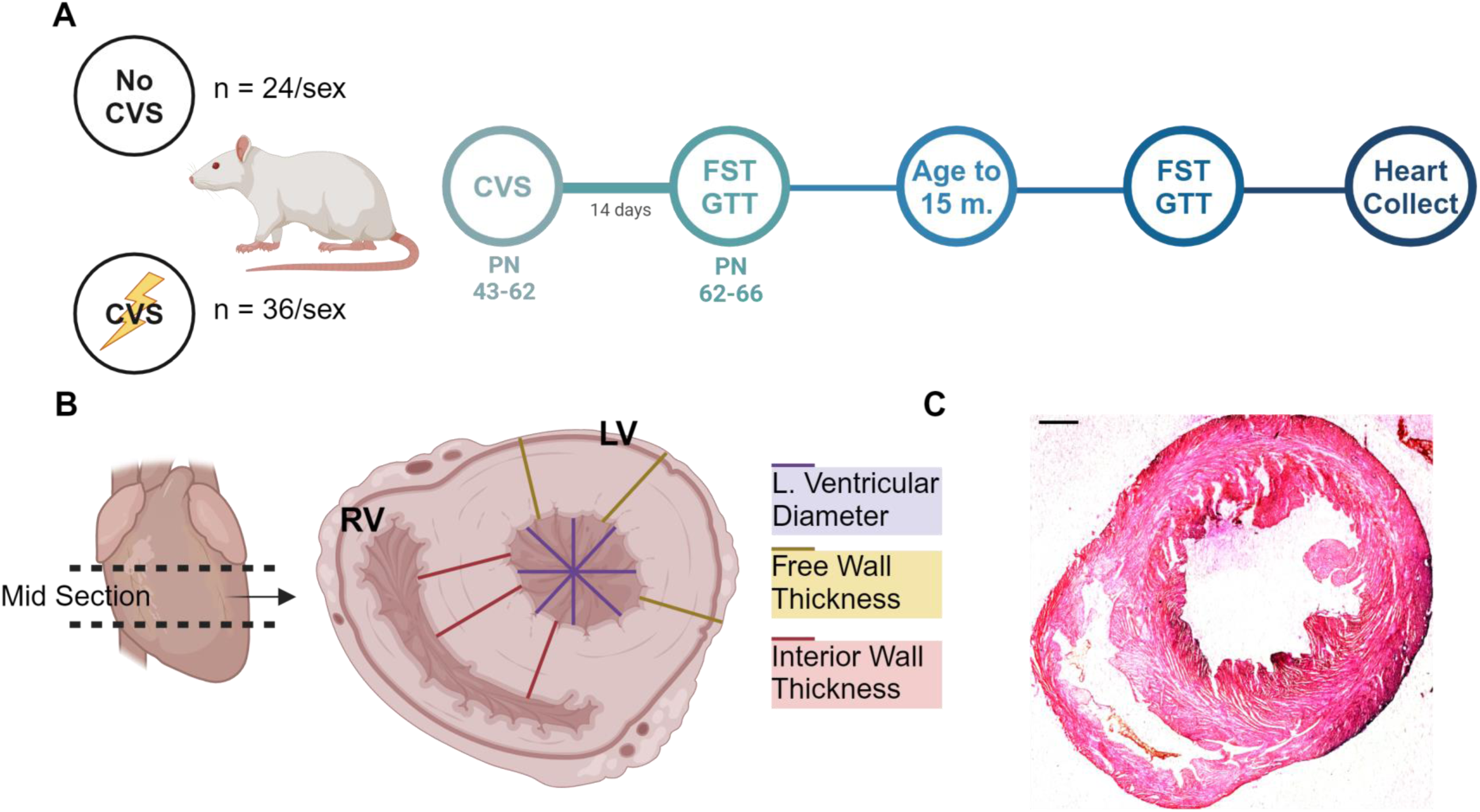
Experimental design. A single cohort (N = 120) of male and female rats underwent chronic variable stress (No CVS n = 24/sex; CVS n = 36/sex) followed by acute challenge and aging (A). Mid-sections of each heart were stained with H&E and wall and ventricular thickness were measured (B,C). Created with BioRender.

### 2.3 Cardiac Tissue Collection

Two days after the final GTT, rats were anesthetized (5% inhaled isoflurane) and euthanized via rapid decapitation. Hearts were immediately arrested in diastole via intracardiac injection of potassium chloride (15 %) followed by saline retroperfusion to remove blood. Heart tissue was then placed in 4% paraformaldehyde for 24 hours and transferred to 30% sucrose for storage. All somatic and organ measures were corrected for bodyweight at the time of euthanasia.

### 2.4 Histologic Processing

Prior to histological evaluation, cardiac tissue was dehydrated, embedded in paraffin, and sectioned. The heart was blocked horizontally into 3 increments to isolate the midsection (Fig. 1B) and placed in a cassette for dehydration. Tissue was incubated in 60% ethanol for 10 minutes and transferred to 70% ethanol for two 20-minute increments in different baths. The tissue was then soaked in two 95% ethanol baths for 20 minutes and transferred to 100% ethanol for three 20-minute incubations. Tissue was transferred to 100% xylene for a total of 90 minutes. After dehydrating the heart tissue, cassettes were dried and placed in paraffin overnight in a Shandon Histocentre 3 (Thermo Electron Cooperation, Waltham, MA). Heart tissue was sectioned using a Microm HM 330 microtome (Microm, Whitney, United Kingdom) at 35 μm. Slices were stained with Hematoxylin and Eosin (Fig. 1C) before drying, cover-slipping, and imaging.

### 2.5 Imaging and Analysis

Brightfield microscopy was performed using a Ziess Axio Imager Z2 microscope (Carl Ziess AG, Oberkochen, Germany) with 9 tissue sections imaged per animal. FIJI Image processing (National Institutes of Health, Bethesda, MA) was used to quantify images with three parameters measured: left ventricle diameter, left ventricular free wall thickness, and interior wall thickness. To measure free wall and interior wall, 3-6 measurements were made per image. The interior wall was measured from the endocardium of the left ventricle wall to the endocardium of the right ventricle. The free wall was measured from the endocardium of the left ventricle to the serosa. The ventricular diameter was measured at 0, 45, -45, and 90 degrees, spanning the entirety of the ventricle. All parameter measurements were then averaged across slices for each animal (n = 36-54 biologic replicates/animal).

### 2.6 Data Analysis

Data are expressed as mean ± standard error of the mean. All rats were included in analyses. Data were analyzed using Prism 10 (GraphPad, San Diego, CA), with statistical significance set at p < 0.05 for all tests. Heart size and VH ratio were analyzed by 2-way ANOVA with sex and stress as factors. Comparison of VH within-group was analyzed by 3-way ANOVA with sex, stress, and hypertrophy as factors. All other comparisons of behavioral, somatic, and physiologic responses to FST and GTT were analyzed by 3-way ANOVA with sex, stress, and hypertrophy as factors. Tukey multiple comparisons post-tests were used when significant main or interaction effects were present. Area under the curve (AUC) analysis was calculated as previously reported [21]. Effect size is reported as eta squared (^2^). Pearson correlation two-tailed analyses were used to examine within-group relationships between VH and homeostatic parameters.

## 3. Results

The current data focus on inward left ventricular hypertrophy and associated factors. Our prior work reports group measures from the FST and GTT [21]. Therefore, while main effects are stated below, sex- and stress-specific differences are outlined in the supplemental material.

### 3.1 Heart Size and Hypertrophy

Heart size was determined by 3 measurements: left ventricular diameter, interior wall thickness, and free wall thickness (Fig. 2). Two-way ANOVA comparison of ventricular diameter found a main effect of stress [F(1, 111) = 11.62, p = 0.0009, ^2^ = 9.326] with post-hoc analysis indicating the No CVS females had larger ventricle diameter than No CVS males (Fig. 2A; p = 0.0437). Interior (Fig. 2B) and free (Fig. 2C) wall measurements both exhibited main effects of sex [F(1, 110) = 13.71, p = 0.0003, ^2^ = 11.0] and [F(1, 111) = 19.08, p < 0.0001, η = 14.58], respectively. Post-hoc analysis of the interior wall measurements indicated CVS females had thicker interior walls than CVS males (p = 0.0144). This was reflected in the free wall analysis with CVS females (p = 0.0082) and No CVS females (p = 0.0177) having significantly thicker free walls than their male counterparts. There were no significant differences within-sex for ventricle measurements.

**Fig. 2.**
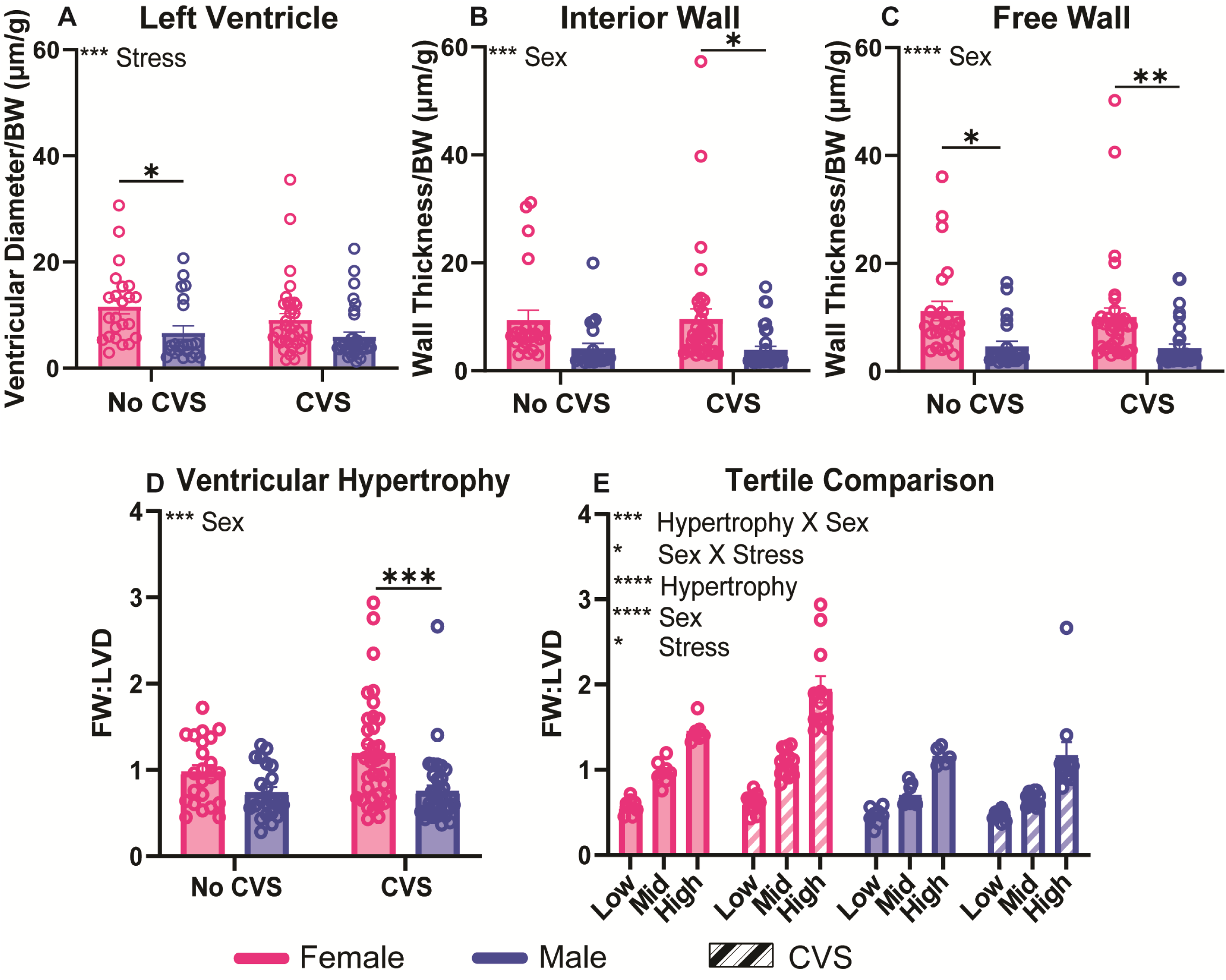
Heart and hypertrophy measurements. The left ventricle (A), interior wall (B), and free wall (C) were measured and corrected for bodyweight at the time of tissue collection. Ventricular hypertrophy (VH) was represented by a ratio of the free wall to the left ventricular diameter (D). Each group was then separated into subgroups (n = 8/sex No CVS and n = 12/sex CVS each for low, mid, and high), and these were compared to determine differences between subpopulations (E). Data are expressed as mean ± SEM. * p<0.05, ** p<0.01, *** p<0.001, **** p<0.0001.

Concentric left VH was determined by a ratio of the free wall thickness to the left ventricular diameter (Fig. 2D). Analysis indicated a main effect of sex [F(1, 111) = 14.39, p = 0.0002, ^2^ = 11.0] where CVS females had increased inward remodeling compared to CVS males (p = 0.0009). An internal comparison of tertiles (n = 8/sex No CVS and n = 12/sex CVS) representing low, mid, and high VH within each sex and stress condition (Fig. 2E) indicated main effects of hypertrophy [F(2, 103) = 118.1, p < 0.0001, ^2^ = 50.77], sex [F(1, 103) = 49.89, p < 0.0001, η^2^ = 10.72], and stress [F(1, 103) = 4.717, p = 0.0322, ^2^ = 1.014]. Additionally, interactions of hypertrophy X sex [F(2,103) = 5.932, p = 0.0036, ^2^ = 2.550] and sex X stress [F(1, 103) = 5.520, p = 0.0207, ^2^ = 1.186] were present with the high hypertrophy group greater than mid and low in all conditions (Table 1). All within-sex and -stress subgroup comparisons were significant except for mid vs low in males and No CVS females (Fig. 2E). Overall, these data indicate that females had larger body weight-corrected heart size that culminated in increased inward hypertrophic remodeling after CVS. Further, the distribution of individuals into subgroups based on VH indicated significant population differences in VH that were moderated by sex, stress, and sex x stress interactions.

**Table 1:**
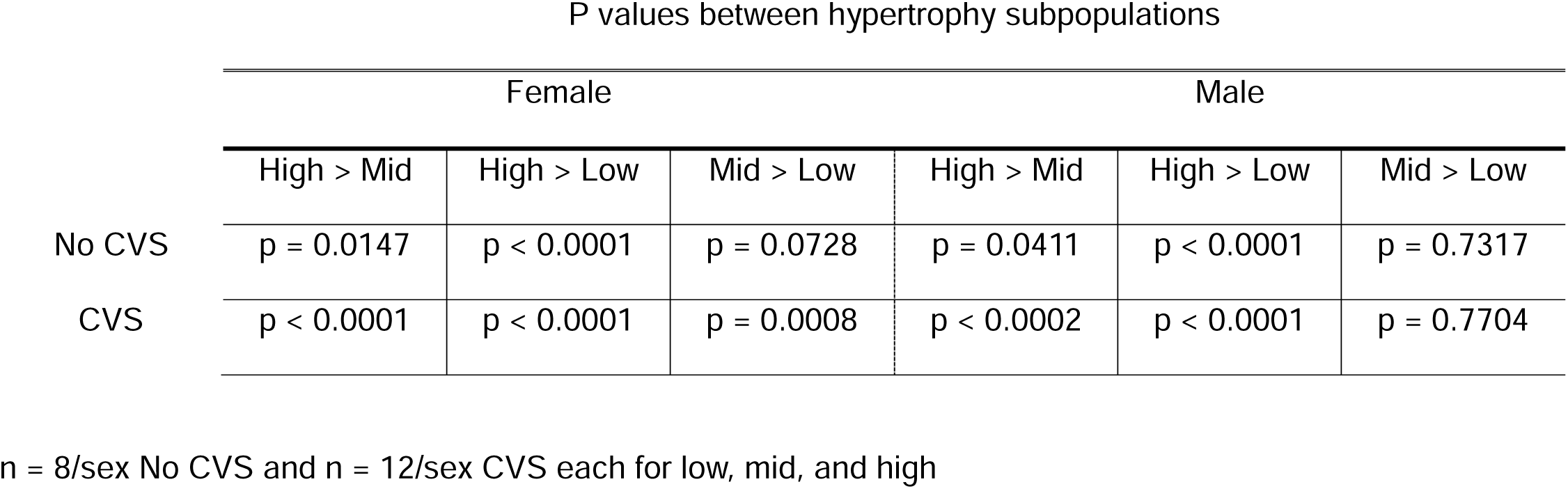
Statistical comparison of hypertrophy subpopulations by group.

### 3.2 Coping Behaviors

Behavioral coping style (Fig. 3) was impacted by subsequent VH depending on the age of the animals. When separated into VH subpopulations, young passive coping (Fig. 3A), represented by immobility during FST immediately following CVS, had main effects of sex [F(1,103) = 31.18, p < 0.0001, η^2^ = 20.26], stress [F(1, 103) = 4.641, p = 0.0335, η^2^ = 3.015] and future VH [F(2, 103) = 4.786, p = 0.0103, η^2^ = 6.218] with an interaction between VH and stress [F(2, 103) = 3.148, p = 0.0471, ^2^ = 4.09]. Young active coping, represented by swimming during FST, was not significantly impacted by VH (Fig. 3B). However, analysis found main effects of sex [F(1, 103) = 17.82, p < 0.0001, η^2^ =12.88] and stress [F(1, 103) = 10.04, p = 0.002, ^2^ = 7.258], with the VH-susceptible CVS males showing less active coping than No CVS counterparts (p = 0.0462).

**Fig. 3.**
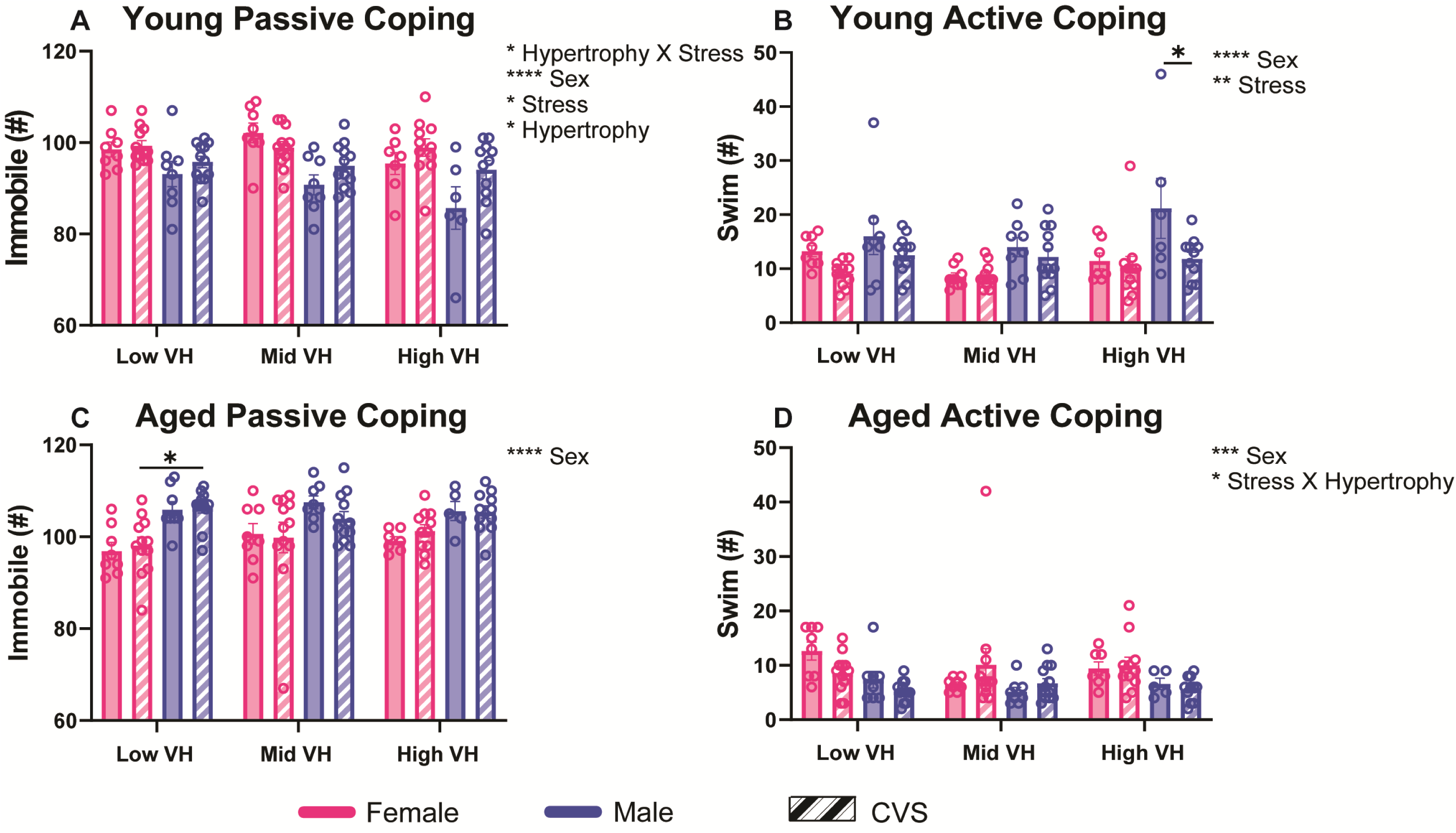
Coping behavior. Animals were challenged with a forced swim test immediately following chronic variable stress and after aging. Coping style was assessed as passive (A, C) or active (B, D). These were then compared across hypertrophy subpopulations (n = 8/sex No CVS and n = 12/sex CVS each for low, mid, and high). Data are expressed as mean ± SEM. * p<0.05, ** p<0.01, *** p<0.001, **** p<0.0001.

In aged animals, immobility (Fig. 3C) was not impacted by VH but maintained a main sex effect [F(1, 102) = 30.75, p < 0.0001, ^2^ = 22.15]. However, within the VH-resilient animals, aged CVS males had more passive coping than CVS females (p = 0.0467). Active coping in aged animals (Fig. 3D) was affected by sex [F(1, 102) = 14.52, p = 0.0002, ^2^ = 11.36] and the interaction between stress exposure and hypertrophy [F(2, 102) = 3.565, p = 0.0319, ^2^ = 5.583]. Hormonal responses to FST (Fig. S1) did not have main effects or interactions that involved hypertrophy. Instead, the sex and stress effects previously reported for CVS to modify glucose mobilization and corticosterone secretion in a sex-specific manner [21] largely persisted after tertile separation. Taken together, stress coping behaviors across the lifespan were mediated by sex, stress history, VH susceptibility, and interactions between CVS and VH. Specific subgroup differences indicate that the young CVS-exposed males most susceptible to VH had reduced active coping compared to controls. Alternatively, the aged CVS-exposed females least susceptible to VH had less passive coping than chronically-stressed males.

### 3.3 Glucocorticoid Responses to Metabolic Stress

Glucose clearance and glucocorticoid responses during hyperglycemia were not impacted by VH in young rats (Fig. S2). However, after aging, baseline corticosterone and glucocorticoid responses to metabolic stress were impacted by VH in a sex-dependent manner (Fig. 4). Baseline corticosterone (Fig. 4A) in aged animals was affected by sex [F(1, 85) = 4.912, p = 0.0293, ^2^ = 0.0293] and an interaction of sex, stress, and VH [F(2, 85) = 3.283, p = 0.0423, ^2^ = 0.0423]. Moreover, peak corticosterone levels 30 minutes after intraperitoneal glucose injection (Fig. 4B) had a main effect of sex [F(1, 86) = 6.772, p = 0.0109, ^2^ = 6.818] and total glucocorticoid release after hyperglycemia (Fig. 4C) was impacted by an interaction of VH and sex [F(2, 35) = 5.042, p = 0.0119, ^2^ = 20.66]. There were no significant differences in cumulative glucose clearance (Fig. 4D). Given the complex interaction between sex, stress, and VH on baseline corticosterone, additional regressive analysis is described below to isolate individual associations.

**Fig. 4.**
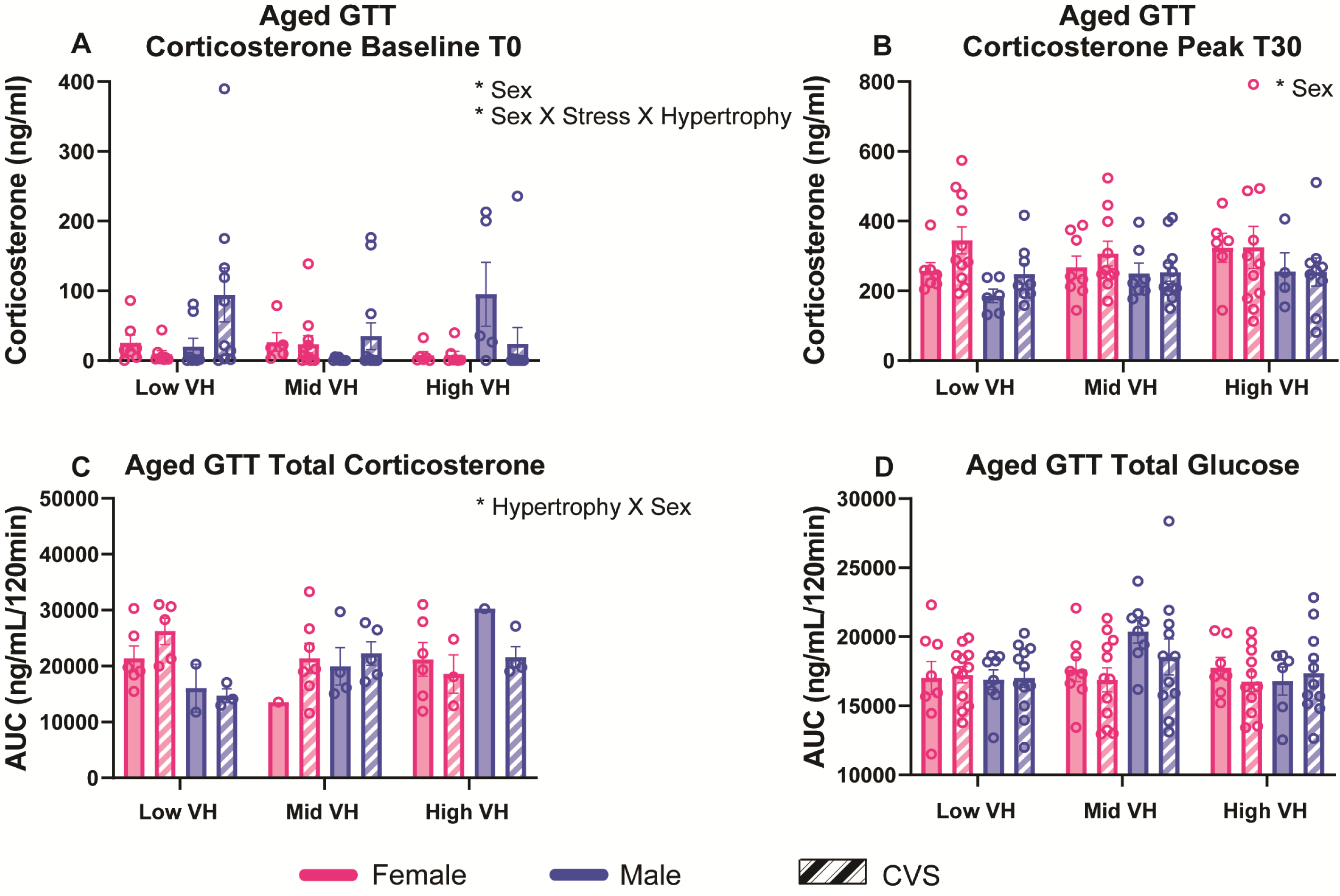
Glucose tolerance. Aged animals were metabolically challenged following chronic variable stress. Baseline corticosterone was measured at 0 min (T0) taken prior to intraperitoneal glucose injection (A). Peak corticosterone was measured 30 minutes following glucose injection (B). Total plasma corticosterone (C) and blood glucose (D) were calculated from an AUC analysis. Groups were analyzed according to hypertrophy subpopulations (n = 8/sex No CVS and n = 12/sex CVS each for low, mid, and high). Data are expressed as mean ± SEM. * p<0.05.

### 3.4 Adiposity

Measures of both visceral (mesenteric) and subcutaneous (inguinal) adiposity were significantly impacted by VH. Mesenteric adiposity (Fig. 5A) had a main effect of VH [F(2, 101) = 3.916, p = 0.023, ^2^ = 6.755]. Moreover, inguinal adiposity (Fig. 5B) was affected by sex [F(1, 100) = 94.98, p < 0.0001, ^2^ = 41.97] and an interaction between VH, sex, and stress [F(2, 100) = 3.09, p = 0.499, ^2^ = 2.731]. Among the VH-resilient groups, CVS males had greater inguinal adiposity than CVS females (p = 0.0038). Additionally, male rats in the mid VH group had greater inguinal adiposity than females regardless of stress (No CVS: p < 0.0001; CVS: p = 0.0005). In the VH-susceptible groups, CVS males had higher adiposity than CVS females (p < 0.0001). Additional somatic measures (Fig. S3) identified primary sex effects that did not relate to VH across spleen and adrenal indices, plasma triglycerides and cholesterol immediately following CVS, as well as body weight after CVS and aging. Collectively, subcutaneous adiposity was regulated by interactions between sex, stress history, and VH where chronically-stressed males had greater subcutaneous adiposity than CVS females across risk/resilience subpopulations.

**Fig. 5.**
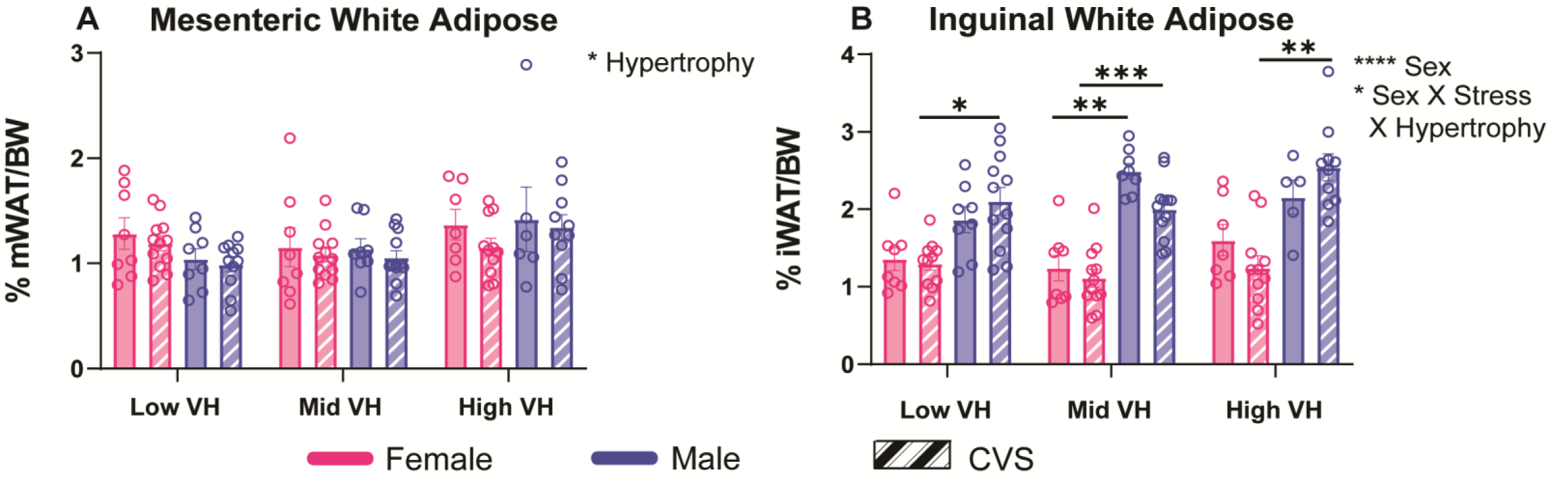
Adiposity. Following aging, mesenteric white adipose (A) and inguinal white adipose (B) tissues were measured. Groups were analyzed according to hypertrophy subpopulations (n = 8/sex No CVS and n = 12/sex CVS each for low, mid, and high). Data are expressed as mean ± SEM. * p<0.05, ** p<0.01, *** p<0.001, **** p<0.0001.

### 3.5 Correlative Analyses

In the cases where ANOVA found main or interaction effects with VH, regressive analyses were performed within stress (No CVS/CVS) and sex (female/male) conditions to examine individual associations with VH susceptibility. This approach revealed that VH positively correlated with baseline corticosterone only in aged No CVS male rats (r = 0.449, p = 0.041; Fig. 6A). Furthermore, VH positively correlated with visceral adiposity (r = 0.387, p = 0.026) only in aged males that were previously exposed to CVS (Fig. 6B). Interestingly, there were no significant correlations with VH in females across any measure, regardless of stress history. Ultimately, aged males specifically had endocrine and metabolic associations with VH susceptibility that related to early-life stress history.

**Fig. 6.**
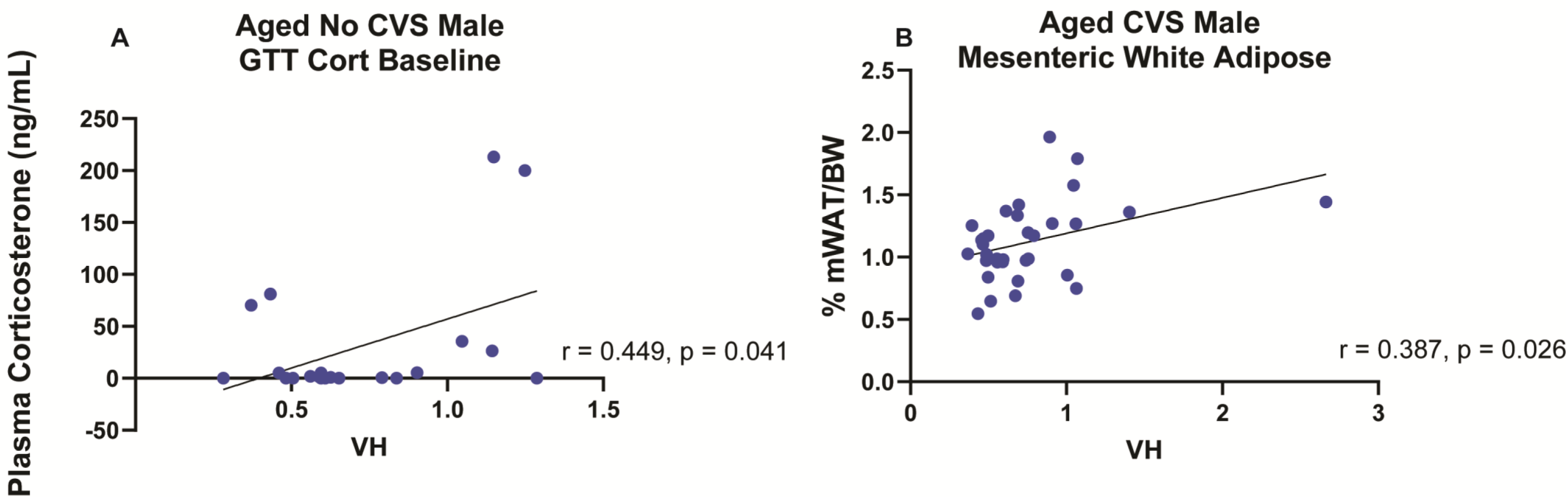
Correlations. Regressive analysis (No CVS n = 24/sex; CVS n = 36/sex) identified VH associations of aged No CVS male baseline corticosterone (A) and aged CVS male visceral adiposity (B).

## 4. Discussion

### 4.1 Female-Specific Cardiac Hypertrophy

The current study sought to examine the effects of stress on cardiac structure after aging in male and female rats. While early-life chronic stress had a modulatory influence on cardiac structure later in life, the primary multi-system effects were driven by sex differences across the aging process. Heart size and VH ratio indicated that females had significant differences in relative cardiac structure. However, these changes led to deleterious remodeling after early-life CVS in females specifically. This contrasts with prior reports of chronic stress-induced cardiac hypertrophy in young male, but not female, rats [23]. These age-specific results are in accordance with clinical reports that rates of CVD increase in women relative to men with aging [3]. While the mechanisms behind these changes in risk are not completely understood, there is an emerging consensus that female reproductive hormones offer cardiovascular protection [5,6]. Thus, female susceptibility in our study may have continued to increase if aging had extended beyond reproductive senescence. Importantly, inward ventricular remodeling in females exposed to early-life chronic stress indicates that early adversity impacts cardiovascular structure across the lifespan in a sex-specific manner that is not entirely dependent on the loss of reproductive hormones. These results may help to explain the finding that female CVD rates rapidly overtake and exceed those of males following menopause [4], as females may be predisposed to structural changes that are masked prior to menopause.

### 4.2 Male Variation in Susceptibility

Exposure to CVS causes male-specific cardiac remodeling immediately following stress exposure [24]. However, this effect seems to be eclipsed by aging. While aged chronically-stressed males did not experience VH as a group, individuals with the greatest VH exhibited homeostatic dysfunction in a stress history-specific manner. This susceptibility indicates that the long-term deleterious effects of early-life chronic stress exposure may differentially impact individuals of the same population. Additionally, susceptible individuals had different behavioral coping responses to acute stress that were related to later measures of VH. Specifically, the interaction of VH and stress within the young passive coping response indicates that, in males, later cardiac remodeling is predicted by behavioral responses to acute challenge depending on cumulative stress burden. Further, unstressed males with the greatest VH had increased active coping compared to their CVS counterparts. The increase in active coping may be an adaptive response to acute stress in naïve animals that then increases susceptibility to cardiac remodeling. Chronic stress exposure appears to dampen these changes and adaptive responses. These results indicate that, in males, early-life stress and consequent coping behavior predict later cardiac changes. Interestingly, passive coping later in life does not correlate with cardiac remodeling and is instead primarily a function of sex.

While endocrine responses to the FST did not identify male susceptibility in either young or aged animals, acute metabolic stress in aged animals emphasized stress- and sex-specific susceptibility in males. Particularly, baseline corticosterone measurements in aged animals had an inverse relationship between stress condition and VH. Additionally, regressive analysis found a positive correlation between VH and baseline corticosterone in unstressed males only. These results suggest that increased glucocorticoid exposure in unstressed aged males associates with greater inward hypertrophic remodeling, a relationship that is disrupted by chronic stress. Further, the positive correlation in unstressed males implicates a deleterious association between glucocorticoid tone and cardiac remodeling that becomes apparent with age.

The association between metabolic processes and aged cardiac hypertrophy is also apparent in measures of adiposity. Visceral adiposity is strongly correlated with the development of multiple cardiometabolic diseases in humans [25]. The correlation between subcutaneous adiposity and CVD is less clear. However, an overall increase in adiposity is persistently associated with increased risk for cardiometabolic dysfunction [26]. Interestingly, exposure to chronic stress leads to a positive correlation of visceral adiposity and VH in males, indicating that early-life chronic stress exposure may impact metabolic capacity and consequently cardiac remodeling. The interaction of sex, stress, and VH risk on subcutaneous adiposity further indicates overall adiposity as a mediator of male VH. Interestingly, more traditional measures of cardiovascular risk, triglycerides and cholesterol [27], were not associated with VH, although both measures were significantly impacted by sex. However, these measures were taken in young animals immediately following chronic stress exposure. While post-stress measures of these metabolic factors did not predict later VH, they do not incorporate the effects of aging.

### 4.3 Bimodal Dysfunction

Exposure to chronic stress and consequent repeated activation of the HPA axis and autonomic nervous systems impact cardiac workload [28]. In some animals, particularly chronically-stressed females, this led to deleterious inward hypertrophy of the left ventricle. However, the subpopulations of animals with low VH may also harbor maladaptation. This may account for outcomes where both low VH and high VH subpopulations similarly deviate from the mid VH groups. For instance, among unstressed aged female, both the low and mid VH subpopulations have greater glucocorticoid reactivity to hyperglycemia than mid VH females. This pattern of stress- and sex-specific bimodal distributions occurs in multiple homeostatic measures and may indicate maladaptive consequences for deviations in VH. Comparable bimodal responses have been reported elsewhere including findings that rats with low glucocorticoid responsiveness following stress have similar sympathetic dysregulation to highly-responsive animals [29].

## 5. Conclusions

The results of the current study indicate that adolescent chronic stress exposure sex-specifically impacts cardiometabolic physiology immediately following stress. However, the impact of aging unmasks a female-specific vulnerability to cardiac hypertrophy. Moreover, in male rats, behavioral, metabolic, and endocrine responses to acute stress associate with cardiac hypertrophy, indicating the potential to identify susceptible subpopulations. Importantly, these results suggest that sex-specific responses following chronic stress differentially impact later individual cardiac health. Thus, while some individuals showed resilience, those that were susceptible to VH may be at increased risk for CVD. Ultimately, defining the sex-specific factors that mediate the impact of early-life adversity on later CVD risk could led to improved approaches for personalized care.

## 7. Declarations

### 7.1 Ethics Approval and Consent to Participate

Animal experiments were approved by the Institutional Animal Care and Use Committee of the University of Cincinnati (protocol 04-08-03-01) and complied with the National Institutes of Health Guidelines for the Care and Use of Laboratory Animals.

### 7.2 Consent for Publication

Not applicable

### 7.3 Availability of Data and Materials

The cohort data were previously published [21] https://doi.org/10.1016/j.yhbeh.2021.105060. All data generated during the current study are available upon request.

### 7.4 Competing Interests

The authors declare that they have no competing interests.

### 7.5 Funding

This research was supported by NIH grants F30 OD032120 to C. Dearing and R01 HL150559 to B. Myers, as well as a Pilot Translation Research Program grant to B. Myers and L. Wulsin from the University of Cincinnati Department of Psychiatry and Behavioral Neuroscience. N. Olmstead was supported by NIH training grant T35 OD015130 (PI: Zabel).

### 7.6 Authors Contributions

C. Dearing, R. Morano, L. Wulsin, and B. Myers designed the research; C. Dearing, E. Sanford, N. Olmstead, R. Morano, and B. Myers performed the experiments; C. Dearing and E. Sanford analyzed the data; C. Dearing, E. Sanford and B. Myers wrote the manuscript; all authors edited the manuscript and approved the final submitted version.

## 7.7 Acknowledgements

The authors would like to thank Elaine Ptaskiewicz for behavioral analysis, Benjamin Schneider for contributions to histology protocol development, and Tyler Wallace for data management. We are grateful for the overall experimental support of Parinaz Mahbod, Jessie Scheimann, and Ana Franco-Villanueva. The authors would also like to thank the many colleagues of the University of Cincinnati Reading Campus that contributed to sample and tissue collection.

## Supplemental Material

### Endocrine Responses: psychogenic stress and metabolic stress

Analysis of forced swim test (FST) responses showed numerous sex and stress effects (Fig. S1). Baseline corticosterone measures indicated main effects of sex [(1, 103) = 8.650, p = 0.004, ^2^ = 7.006] and stress [F(1, 103) = 4.706, p = 0.0324, ^2^ = 3.812] in young animals and a main effect of sex [F(1, 102) = 24.03, p < 0.0001, η = 17.7] in aged animals. This is maintained in the young total corticosterone response with both main effects of sex [F(1, 38) = 13.66, p = 0.0007, η^2^ = 14.78] and stress [F(1, 38) = 27.81, p < 0.0001, ^2^ = 30.09]. However, aged total corticosterone response to FST shows no significant effect. Significant sex differences were noted in both the young and aged total blood glucose response to FST with a main effect of sex [F(1, 101) = 48.31, p < 0.0001, ^2^ = 29.31] and [F(1, 102) = 67.10, p < 0.0001, ^2^ = 37.81], respectively. Interestingly, within the young mid VH group, young No CVS females mounted a smaller glucose response than both CVS females (p = 0.0253) and No CVS males (p = 0.0007). Within the high VH young animals, CVS males had a greater glucose response than CVS females (p = 0.0047). Aged animals also showed sex-specific differences with the low VH No CVS males having a greater glucose response than No CVS females (p = 0.0076) and CVS males showing a greater glucose response than their female counterparts in both the mid VH (p = 0.0019) and High VH (p = 0.0212) groups.

While glucose tolerance test (GTT) glucocorticoid responses in aged animals were impacted by VH, GTT responses in young animals showed only sex effects (Fig. S2). Young baseline corticosterone measures showed a main sex effect [F(1, 90) = 32.51, p < 0.0001, ^2^ = 24.63]. When looking at peak corticosterone levels 30 minutes following intraperitoneal injection of glucose, young animals show no significant differences. Similarly, young animals show no significant effects in total corticosterone response to metabolic challenge.

**Fig. S1.**
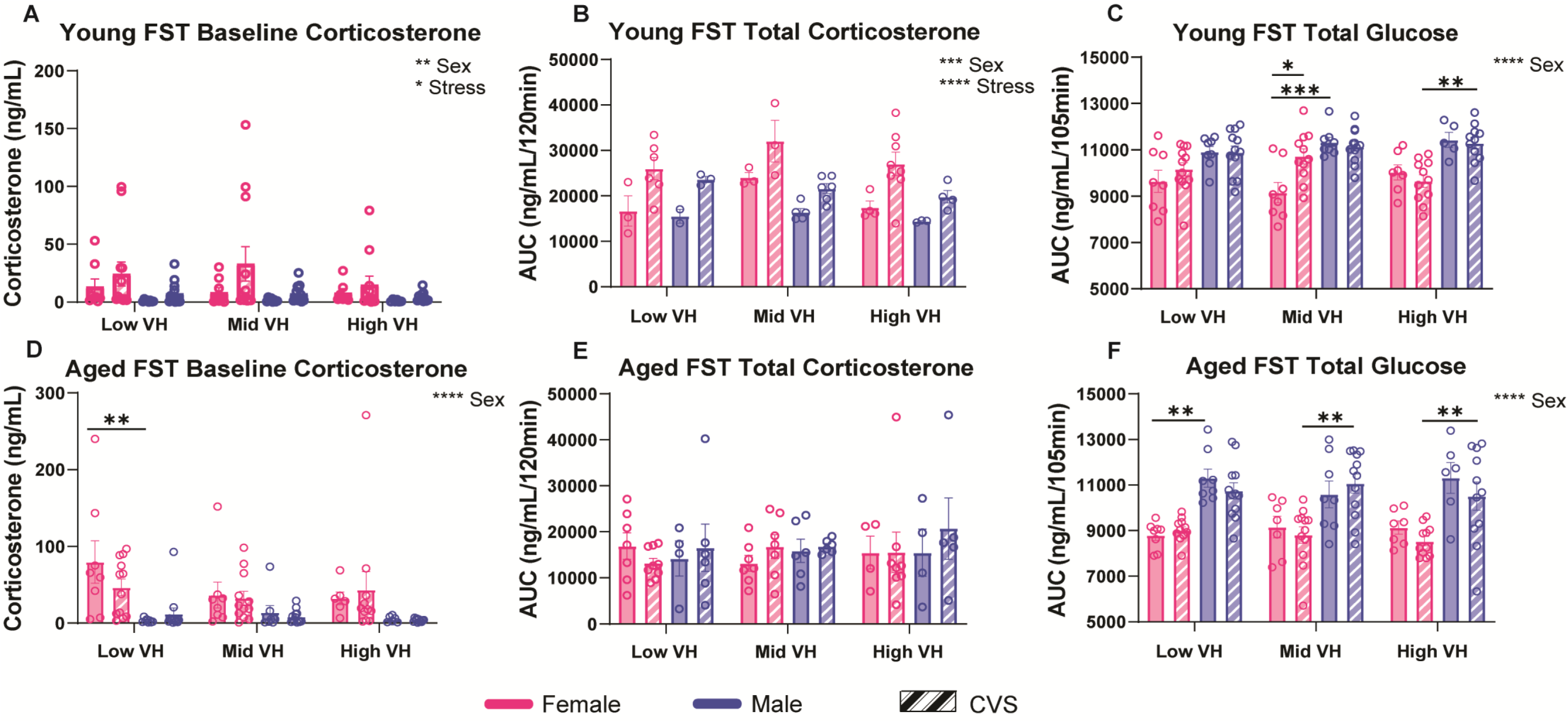
Forced swim test (FST) endocrine analyses. Baseline plasma corticosterone was measured (A,D) two days prior to FST. Total plasma corticosterone and blood glucose were calculated from the AUC for both the young (B,C) and aged (E,F) animals. Groups were analyzed according to hypertrophy subpopulations (n = 8/sex No CVS and n = 12/sex CVS each for low, mid, and high). Data are expressed as mean ± SEM. * p<0.05, ** p<0.01, *** p<0.001, **** p<0.0001.

**Fig. S2.**
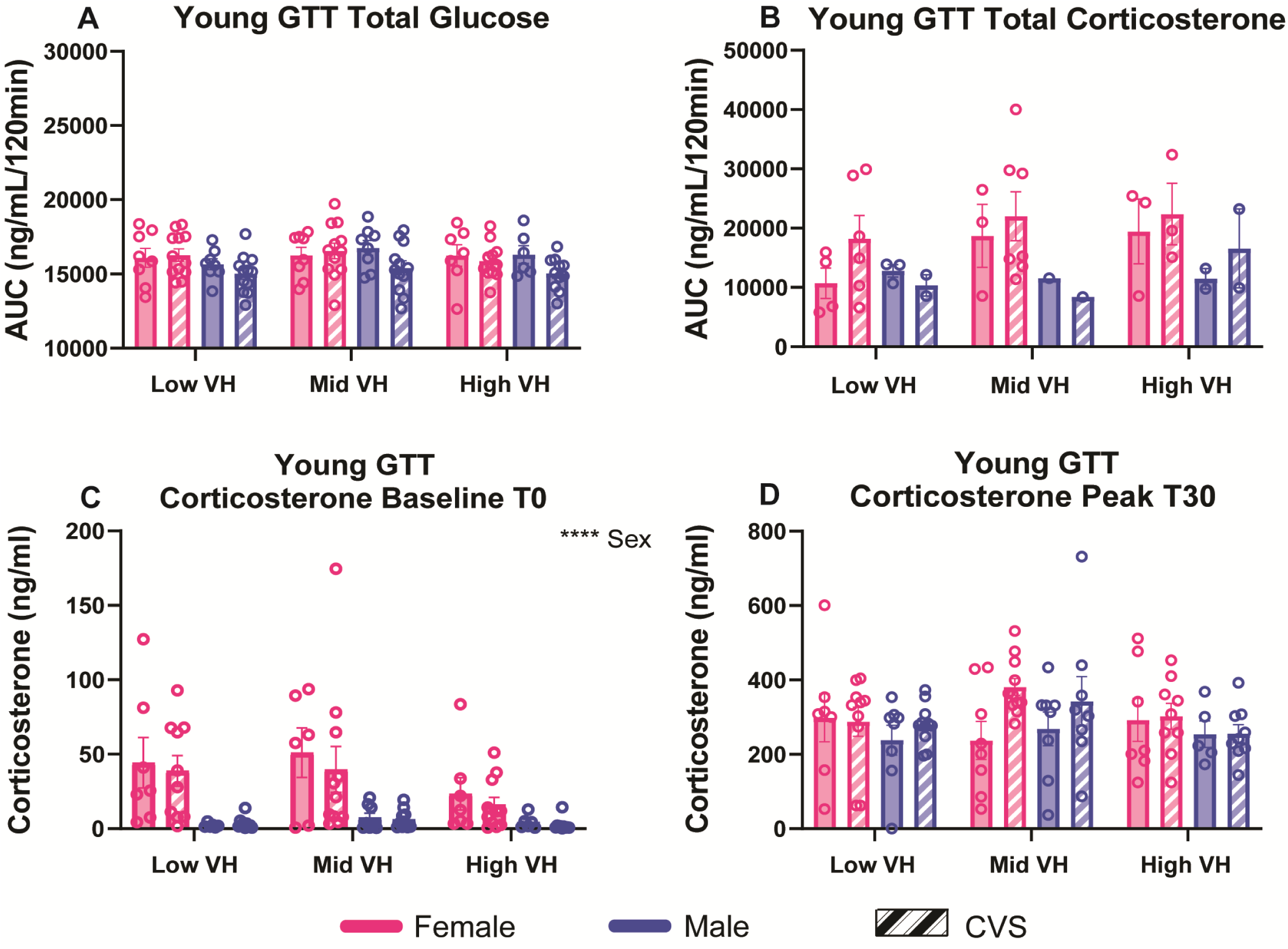
Young glucose tolerance test (GTT) endocrine analyses. Young animals were metabolically challenged following chronic variable stress. Total blood glucose (A) and plasma corticosterone (B) were calculated from an AUC analysis. Baseline corticosterone was measured at T0 taken prior to glucose injection (C). Peak corticosterone response was measured 30 minutes following glucose injection (D). Groups were analyzed according to hypertrophy subpopulations (n = 8/sex No CVS and n = 12/sex CVS each for low, mid, and high). Data are expressed as mean ± SEM. **** p<0.0001.

### Somatic Measures

Somatic measures of body and organ weights showed sex- and stress-specific effects (Fig. S3). Measures of bodyweight-corrected spleen [F(1, 100) = 13.05, p = 0.0005, η^2^ = 10.06] and adrenal weight [F(1, 101) = 230.4, p < 0.0001, η^2^ = 66.19] showed main effects of sex. Main effects of sex were also reflected in young triglyceride levels [F(1, 101) = 84.15, p < 0.0001, η^2^ = 38.71] and plasma cholesterol [F(1, 101) = 16.01, p = 0.0001, η^2^ = 12.36] measured immediately following CVS. Additionally, young triglycerides showed a main interaction of sex and stress [F(1, 101) = 5.10, p = 0.0261, η^2^ = 2.346]. Body weight analysis showed main effects of sex both immediately following chronic stress [F(1, 103) = 819.0, p < 0.0001, η^2^ = 84.04] and at tissue collection [F(1, 103) = 1032, p < 0.0001, η^2^ = 86.23].

**Fig. S3.**
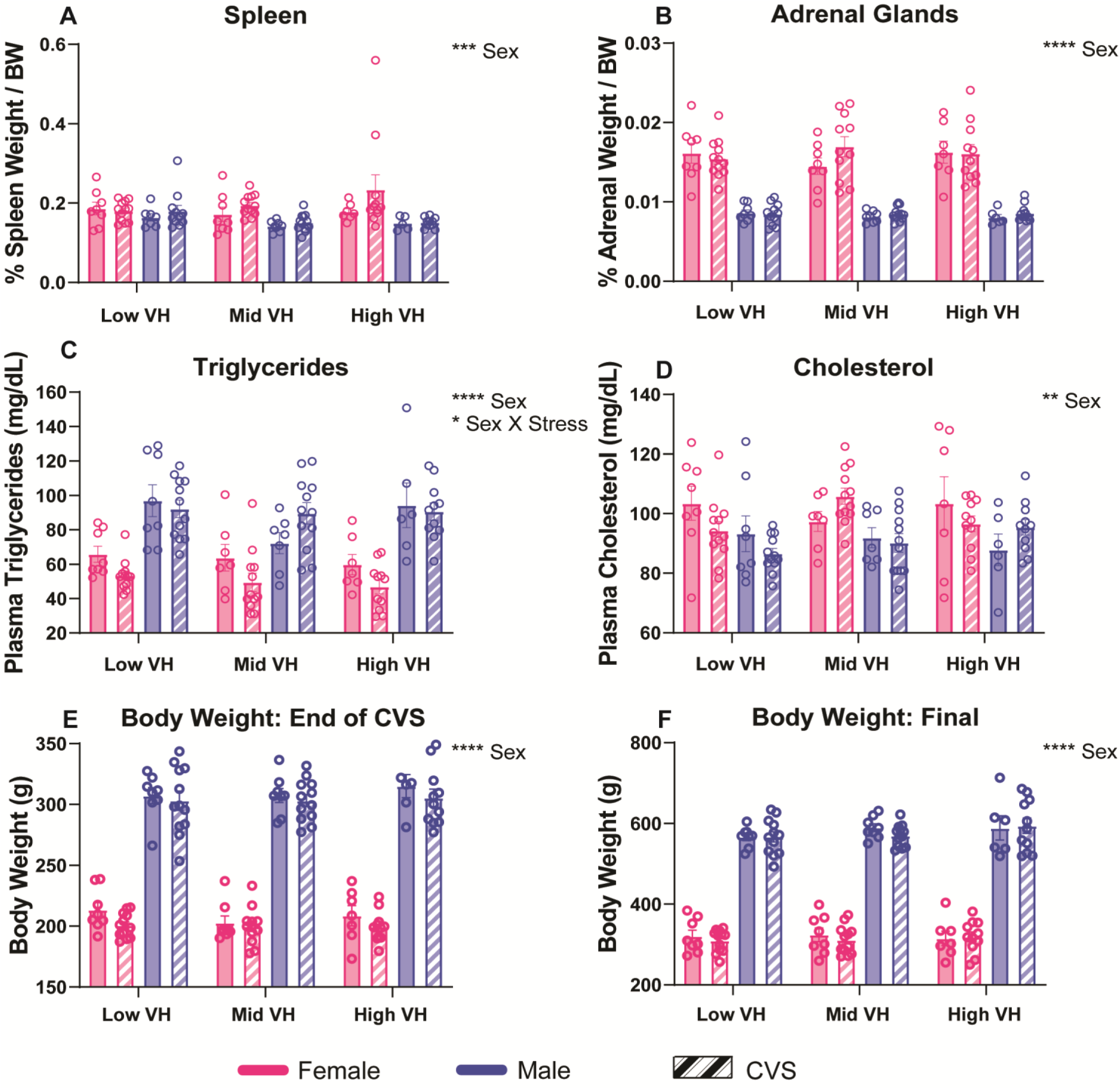
Somatic measures following chronic variable stress. Relative spleen (A) and adrenal weight (B) were measured. Triglycerides (C) and cholesterol (D) were measured following chronic variable stress from baseline blood samples. Body weight was measured and analyzed at both immediately following CVS and at euthanasia (F)). Groups were analyzed according to hypertrophy subpopulations (n = 8/sex No CVS and n = 12/sex CVS each for low, mid, and high). Data are expressed as mean ± SEM. * p<0.05, ** p<0.01, *** p<0.001, **** p<0.0001.

## Notes

### Competing Interest Statement

The authors have declared no competing interest.

